# Assessing cortical excitability with electroencephalography: a pilot study with EEG-iTBS

**DOI:** 10.1101/2023.09.08.556829

**Authors:** Giovanni Pellegrino, Anna-Lisa Schuler, Zhengchen Cai, Daniele Marinazzo, Franca Tecchio, Lorenzo Ricci, Mario Tombini, Vincenzo di Lazzaro, Giovanni Assenza

**Author notes:** **Corresponding Authors** Giovanni Assenza, MD, PhD, Policlinico Universitario Campus Biomedico di Roma, Via Alvaro del Portillo, 200, 00100, Rome, Italy; Giovanni Pellegrino, MD, PhD, Room B10-118, University Hospital, 339 Windermere Road, London, Ontario, Canada.

## Abstract

Cortical excitability measures neural reactivity to stimuli, usually delivered via Transcranial Magnetic Stimulation (TMS). Excitation/inhibition balance (E/I) is the ongoing equilibrium between excitatory and inhibitory activity of neural circuits. According to some studies, E/I could be estimated in-vivo and non-invasively through the modeling of electroencephalography (EEG) signals. Several measures have been proposed (phase consistency in the gamma band, sample entropy, exponent of the power spectral density 1/f curve, E/I index extracted from detrend fluctuation analysis, and alpha power). It remains to be investigated to what extent they scale with excitability and how they relate to each other. Intermittent theta burst stimulation (iTBS) of the primary motor cortex (M1) is a non-invasive neuromodulation technique allowing controlled and focal enhancement of cortical excitability and E/I of the stimulated hemisphere. M1 excitability and several E/I estimates extracted from resting state EEG recordings were assessed before and after iTBS in a cohort of healthy subjects. Enhancement of M1 excitability, as measured through motor-evoked potentials (MEPs), and phase consistency of the cortex in high gamma band correlated with each other. Other measures of E/I showed some expected results, but no correlation with TMS excitability measures or consistency with each other. EEG E/I estimates offer an intriguing opportunity to map cortical excitability non-invasively, with high spatio-temporal resolution and with a stimulus independent approach. While different EEG E/I estimates may reflect the activity of diverse excitatory-inhibitory circuits, spatial phase synchrony in the gamma band is the measure that best captures excitability changes in the primary motor cortex.

## Introduction

Cortical excitability is the predisposition of neuronal populations to generate activity in response to various stimuli (Rossini et al. 2015; Siebner et al. 2022). Cortical excitability is fine tuned across spatial and temporal dimensions, allows proper information integration and is impaired in nearly all neuropsychiatric conditions (Vincenzo Di Lazzaro et al. 2016; 2014; Lazzaro et al. 2017; Landi et al. 2015; Tombini et al. 2013; Di Pino et al. 2014). In the context of human studies, conventional assessment of excitability involves the application of Transcranial Magnetic Stimulation (TMS) and the measurement of elicited behavioural responses, in most cases muscle contractions. TMS pulses interact with neurons spanning all cortical layers, yielding a response shaped by both excitatory and inhibitory neuronal inputs (Vincenzo Di Lazzaro and Ziemann 2013). While TMS has established itself as a potent tool for the study of cortical excitability, it does come with inherent limitations. Firstly, TMS studies primarily focus on eloquent cortices (such as the primary motor cortex or primary visual cortex), where there are relatively easily measurable responses, i.e. muscular contractions or phosphenes. Secondly, when exploring non-eloquent cortices, concurrent neuroimaging measurements are necessary to measure the response to stimuli. Simultaneous TMS-Electroencephalography (TMS-EEG) (Giambattistelli et al. 2014; Tremblay et al. 2019), TMS-functional magnetic resonance imaging (TMS-fMRI) (Tik et al. 2023; Mizutani-Tiebel et al. 2022) and TMS-functional near-infrared spectroscopy (TMS-fNIRS) (Cai et al. 2022) are among the most common options (Bergmann et al. 2016), but the data acquisition process is long and difficult. Neuroimaging signals are affected by artifacts, and there is uncertainty on the extent to which neuroimaging metrics align with the classical definition of cortical excitability (Cai et al. 2022; Conde et al. 2019). Thirdly, whether alone or in combination with neuroimaging, TMS pulses can only be administered at one specific site at a time, leading to a scattered spatial and temporal sampling. Fourthly, TMS pulses recruit a mixture of excitatory and inhibitory neurons in the human cortex (Hannah et al. 2020; Vincenzo Di Lazzaro and Ziemann 2013), and multiple specific TMS protocols are needed to assess specific components of cerebral circuits (i.e. paired-pulse TMS, short-afferent inhibition, silent period, etc.).

In parallel with the development of concurrent TMS-neuroimaging approaches, there has been a remarkable development in the modeling and assessment of the equilibrium between excitatory and inhibitory activity of neural circuits, also known as Excitation/Inhibition balance (E/I). The E/I is linked to the ongoing fluctuations of synaptic input ‘at rest’ and regulate the generation of field potentials (Gao, Peterson, and Voytek 2017).

Currently, modeling of electromagnetic (EEG or MEG) and hemodynamic (fMRI or fNIRS) brain signals are considered reliable approaches for the estimation of E/I (Ahmad et al. 2022; Trakoshis et al. 2020; Zou et al. 2008; Zhang et al. 2010; Cai et al. 2022). For example, in the field of EEG and magnetoencephalography (MEG), E/I may be estimated using EEG phase synchronization (Meisel et al. 2015; 2016; Schuler et al. 2023), aperiodic component of the power spectrum (Gao, Peterson, and Voytek 2017; Donoghue et al. 2020), and entropy (Liang et al. 2015; Wei et al. 2013); in the field of fMRI/fNIRS, E/I can be estimated from the amplitude, regularity, predictability and regional homogeneity of the hemodynamic activity (Zou et al. 2008; Zang et al. 2004; Trakoshis et al. 2020; Cai et al. 2022).

Interestingly, the E/I is one of the strongest determinants of cortical excitability (higher E/I relates to higher excitability), and it can be measured independently from external stimuli (Ahmad et al. 2022; Vincenzo Di Lazzaro and Ziemann 2013), at high spatial and temporal resolution and with better applicability and translational potential than excitability alone (Pellegrino, Machado, et al. 2016; Pellegrino, Hedrich, et al. 2016; 2018; Pellegrino et al. 2020; Chowdhury et al. 2018; Hedrich et al. 2017; Schuler et al. 2022; Machado et al. 2018; Pellegrino, Arcara, Cortese, et al. 2019; Ahmad et al. 2022).

Nevertheless, it remains to be understood if a relationship exists between E/I estimates and cortical excitability as measured from responses to TMS and to what extent different E/I estimates are related to each other.

Here we addressed this topic by performing EEG E/I estimates and TMS excitability measures before and after intermittent theta burst stimulation (iTBS) of the primary motor cortex. iTBS is a non-invasive neuromodulatory technique allowing controlled and focal enhancement of cortical excitability and E/I of the target hemisphere (Huang et al. 2005; V. Di Lazzaro et al. 2011; Suppa et al. 2016; V. Di Lazzaro et al. 2008).

We specifically focused on whether: 1) changes in excitability due to iTBS are linked to changes in E/I estimated with EEG, and 2) whether E/I estimates exhibit an expected modulation as reflected in behavioural outcomes (MEPs).

## Material and methods

We enrolled 11 right-handed healthy adults at Campus Bio-Medico University of Rome. Exclusion criteria were: present/past neuropsychiatric conditions, and medications acting on the central nervous system. All subjects underwent a resting state EEG and primary motor cortex TMS sessions to assess E/I and excitability before and after iTBS of the left primary motor cortex (Figure 1). All participants were asked to abstain from caffeine and alcohol in the two days before the study. This study was approved by the Ethic board of Campus Bio-Medico University of Rome and complied with the Declaration of Helsinki. All participants signed a written informed consent prior to participation.

**Figure 1.**
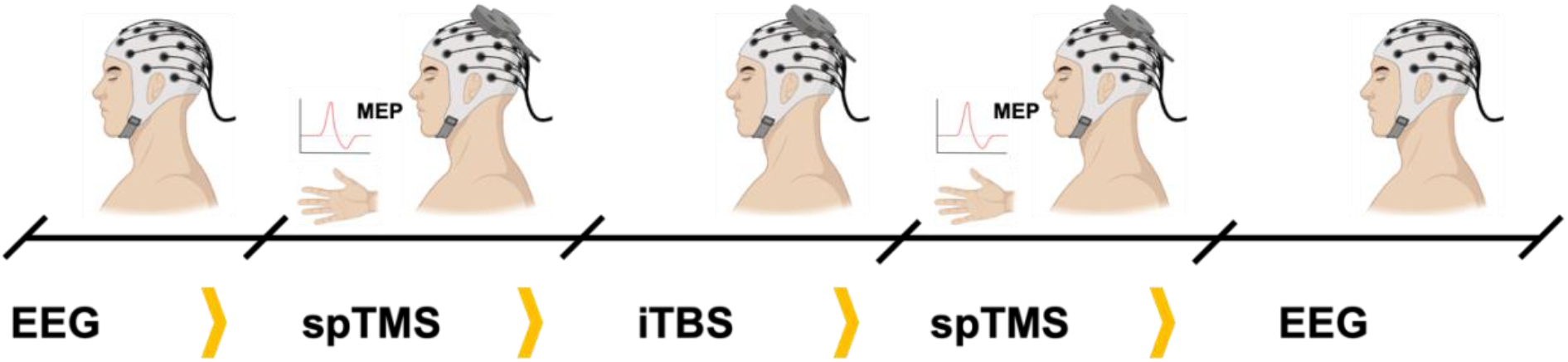
Experimental design: EEG (5 minutes) and spTMS (15 pulses) were acquired before and after iTBS of the left motor cortex. E/I was estimated from EEG, applying several measures, whereas excitability was measured as difference of MEP amplitude before and after iTBS. spTMS: single-pulse TMS, iTBS: intermittent theta burst stimulation. E/I: excitability/inhibition balance.

### EEG recordings

EEG recordings were performed with a BrainAmp 32MRplus amplifier (BrainProducts GmbH, Germany) and 32 electrodes arranged on an elastic cap (https://www.easycap.de), with a sampling rate of 5kHz. The reference was binaural, the ground was placed at the right arm, and electrode impedance was kept below 10 kOhm. EEG data was recorded for 5 minutes in a shielded room, with the subject comfortably lying down and with the eyes open.

### Motor cortex excitability to TMS

Left primary motor cortex excitability was measured by applying 15 single pulses of TMS and recording Motor Evoked Potentials from the right Opponents Pollicis. TMS was performed with a Rapid Magstim Stimulator (Magstim Company, Dyfed, UK) and 70mm eight-shaped coil. MEPs were recorded with surface electrodes positioned according to the standard tendon-belly montage, and connected to the bipolar headset of the BrainAmp 32MRplus amplifier. The Rapid Magstim delivers biphasic stimuli. The hotspot was searched with the TMS coil held approximately over the precentral region, with a 45° direction to the midline. The resting motor threshold was defined as the stimulation intensity able to evoke MEPs in the target muscle >50 microV in 5 out of 10 stimuli (Mioli et al. 2018). The TMS intensity for single pulse TMS was set to 120% of the Resting Motor Threshold.

### iTBS

iTBS was delivered over the left primary motor cortex with standard parameters (Huang et al. 2005; V. Di Lazzaro et al. 2011). The intensity was set to 80% of the active motor threshold (Vincenzo Di Lazzaro et al. 2010). In details, iTBS consisted of short bursts of three TMS pulses delivered at 50Hz, repeated at 5Hz. A sequence of 10 bursts lasted 2 seconds and was repeated every 10 seconds (8 seconds break) for a total of 600 pulses and 190 seconds.

### EEG data analysis

EEG data analysis was performed with the EEGLab toolbox (Delorme and Makeig 2004), the Brainstorm toolbox (Tadel et al. 2011) and custom code in Matlab (The Mathworks, Version 2022b).

For the preprocessing, we first applied a bandpass filter (1Hz – 256Hz), a notch filter, and resampling to 1024Hz. Data were then visually inspected to remove segments affected by artifacts. Channels with excessive noise were interpolated using the spline method. Thereafter, we applied the clean_rawdata (https://github.com/sccn/clean_rawdata) toolbox to remove sudden and brief artifacts and ICA (runica) to separate biological and artifactual EEG components. Artifactual IC component were automatically identified and rejected with the IClabel toolbox (Pion-Tonachini, Kreutz-Delgado, and Makeig 2019), using default parameters. The cleaned EEG was then imported into Brainstorm. We considered a standard MRI template (ICBM 512) (Mazziotta et al. 2001) and standard EEG positions. Sensor data were reconstructed onto the cortical surface with the minimum norm estimate (Hämäläinen and Ilmoniemi 1994). We chose a simple head model with multiple spheres (Pellegrino, Hedrich, et al. 2018). The noise covariance matrix was computed based on the entire EEG signal. After source imaging, estimates of E/I were computed considering the Desikan-Killiani atlas (Desikan et al. 2006; Schuler et al. 2022; 2023; Pellegrino, Arcara, Di Pino, et al. 2019).

### E/I Estimates

We focused on the following E/I estimates: a) Exc_ind; b) exponent of the aperiodic component of the Power Spectrum; c) Sample Entropy; d) Detrended Fluctuation Analysis based E/I index; e) alpha power.

a. ‘Exc_ind’. This index was introduced and validated by Meisel et al. (2015). It corresponds to the mean spatial phase synchronization and, especially in high-gamma frequency bands, correlates well with cortical excitability as estimated via perturbational approaches and modulated via antiseizure medications in epilepsy patients (Meisel et al. 2015; 2016). This E/I estimate roughly corresponds to the well-known phase locking value (PLV), broadly exploited in neuroscience and neurology (Pellegrino, Arcara, Di Pino, et al. 2019; Pellegrino et al. 2022; Pellegrino, Mecarelli, et al. 2018; Pinardi et al. 2023; Bruña, Maestú, and Pereda 2018), computed across space rather than time. We selected the high gamma band (55-95Hz), as recommended in previous studies (Schuler et al. 2023);
b. Exponent of the aperiodic component of the Power Spectrum. This estimate has been introduced recently by Voytek and collaborators (Gao, Peterson, and Voytek 2017; Donoghue et al. 2020). In 2017 Gao et al., provided an elegant demonstration that E/I changes can be estimated from the power law exponent (slope) of the electrophysiological power spectrum (Gao, Peterson, and Voytek 2017). Later on, the same group confirmed this concept with non-invasive and invasive recordings and developed a method to disentangle the periodic vs aperiodic component of the power spectrum and to estimate the exponent of the aperiodic component (FOOOF - fitting oscillations & one over f) (Donoghue et al. 2020). We applied the FOOOF implementation available in the Brainstorm toolbox considering a frequency range between 10Hz and 40Hz (Gerster et al. 2022), and extracted the exponent of the 1/f slope as E/I estimate. A flatter spectrum corresponds to a lower exponent and to higher excitability, whereas a steeper spectrum corresponds to higher exponent and lower excitability (Donoghue et al. 2020).
c. Sample Entropy. Entropy measures have been linked to E/I. Lower entropy is linked to higher E/I, whereas higher entropy is linked to lower E/I (Waschke, Tune, and Obleser, n.d.; Sarasso et al. 2015). Among the many available entropy measures, sample entropy is the most sensitive to E/I variations (Wei et al. 2013; Liang et al. 2015). We computed Sample Entropy by including the code of Cannard (Cannard and Delorme 2022) in the Brainstorm toolbox.
d. Detrend Fluctuation Analysis (DFA) based E/I index. Another index of E/I can be extracted by applying DFA. This approach allows to investigate Long Term Temporal Correlations (LRTC), which are influenced by E/I. Higher values correspond to higher E/I. Recently, it has been demonstrated that this approach is sensitive to E/I alterations known to occur in Alzheimer Disease (Van Nifterick et al. 2023). The computation of DFA-based E/I was performed in alpha band, following the method described in (Van Nifterick et al. 2023) and based on an adaptation for Brainstorm of the code available at https://github.com/annevannifterick/fEI_in_AD;
e. Alpha band power is widely considered a marker of E/I, with higher alpha power corresponding to lower E/I (Pellegrino et al. 2021). Alpha power (8-12 Hz) was computed with the Welch method available in Brainstorm, considering a window length of 1 second, and a window overlap ratio of 50%.

### Statistical approach

Left M1 cortical excitability changes due to iTBS were estimated as POST-PRE MEPs amplitude difference. Similarly, whole cortex changes of E/I balance were computed as the delta (Post-Pre) of all E/I estimates. For some E/I estimates, lower values correspond to higher excitability (see above). In order to have a comparable representation of the effects of iTBS on E/I, we computed the normalized differences, using t-values. POST-PRE was applied for ‘Exc_ind’ and ‘DFA-based E/I’. PRE-POST was used for all other measures. Hence, a positive delta always means an increase of E/I. Based on previous literature (Huang et al. 2005; Suppa et al. 2016; V. Di Lazzaro et al. 2011; 2008) we expect that iTBS increases E/I of the left M1 and left (stimulated) hemisphere. The relationship between excitability (TMS-based) and E/I (EEG-based) of the left M1 (Precentral gyrus of the Desikan Killiany atlas), we computed individual differences of each measure and applied the Spearman’s correlation coefficient.

## Results

*Exc_ind* E/I increased consistently over the bilateral motor and premotor regions (Figure 2, Panel A). Figure 2 Panel B shows the map of Sperman’s correlation between excitability changes estimated from MEPs’ amplitude (Panel C) and individual E/I differences. There is a definite positive correlation between the left sensorimotor region E/I increase and excitability increase (the greater the increase in E/I, the greater the increase in iTBS-related excitability). The relationship is significant in the left M1 (Panel C and D, Rho = 0.627, p=0.019).

**Figure 2.**
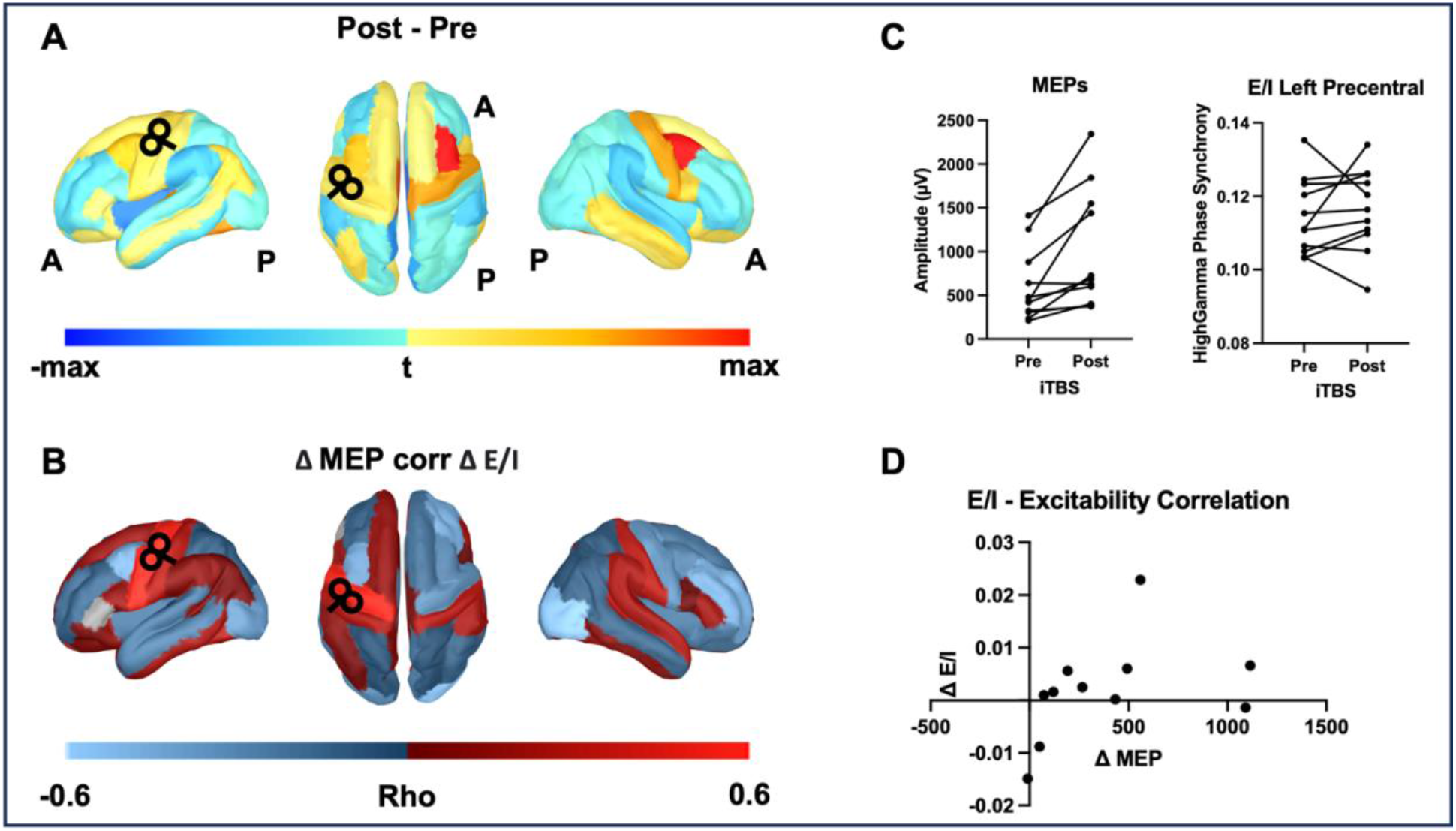
Effects of iTBS on ‘Exc_ind’ extracted from EEG and cortical excitability extracted from TMS-MEPs. *Panel A* shows the normalized Post-iTBS – Pre-iTBS E/I difference. Note the increase in E/I in the bilateral precentral giri, premotor cortex, left primary cortex and bilateral superior frontal cortex. *Panel B* Sperman’s Rho correlation between excitability increase and E/I change, for all regions of the Desikan-Killiany atlas. Note a positive correlation in the left sensorimotor regions. *Panel C Left* Change in MEP amplitude, individual values across subjects. *Panel C Right* Change in E/I, individual values across subjects. *Panel D* Scatterplot of the correlation between E/I and excitability (MEP) changes for the left primary motor cortex (precentral gyrus), where iTBS was delivered (Rho = 0.627, p=0.019).

**Figure 3.**
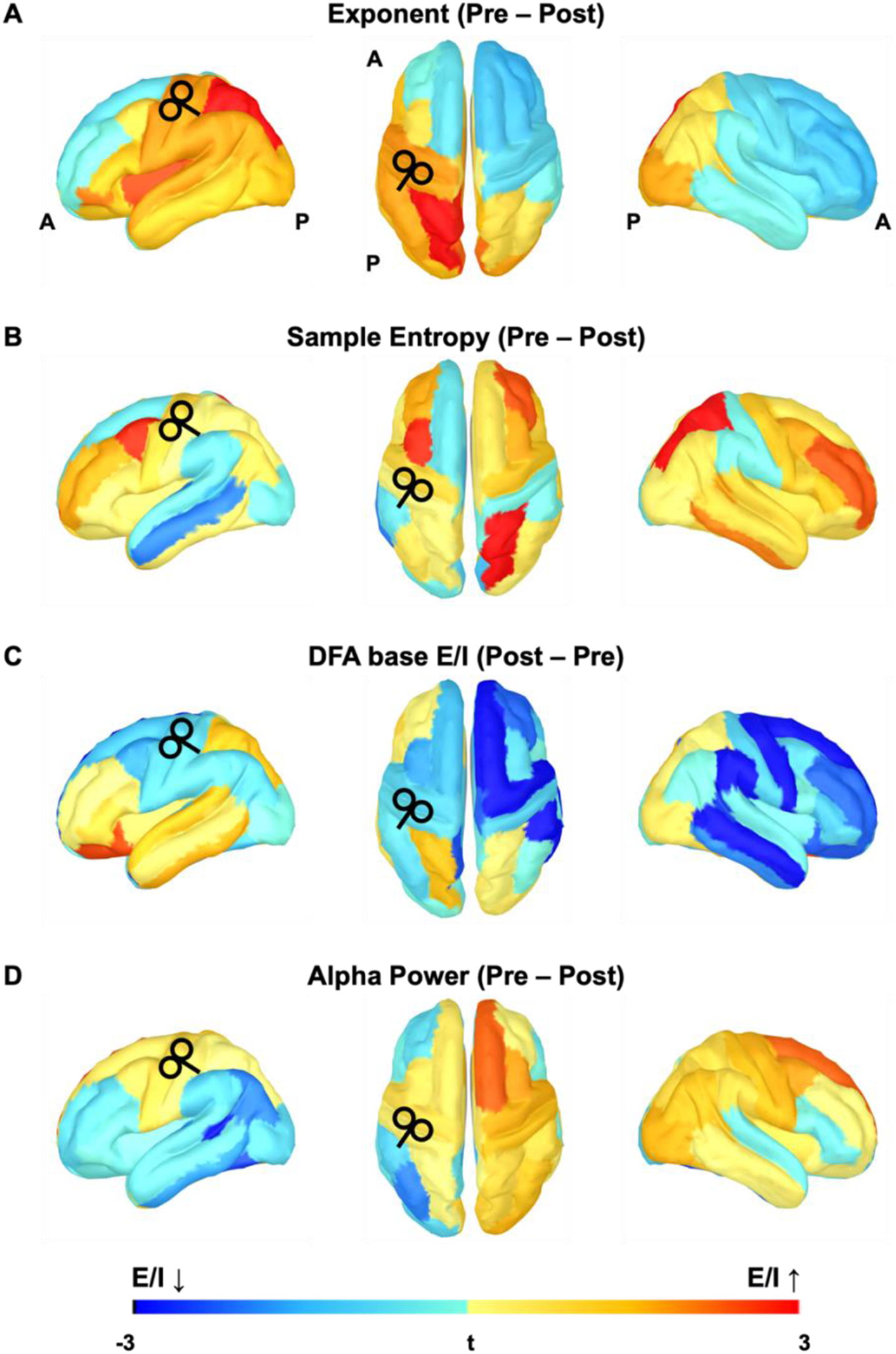
Effects of iTBS on E/I estimates extracted from EEG. Each panel shows the E/I increase induced by iTBS. Panel A: Exponent of the Power Spectral Density; Panel B: Sample Entropy; Panel C DFA-Based E/I; Panel D: Alpha power.

The results for all other E/I estimates are reported in Figure 3. No other E/I index revealed a significant correlation between M1 E/I and excitability changes. The iTBS-related variations show and expected pattern, with increase of E/I in the left hemisphere and especially M1 for the Exponent of the PSD, Sample Entropy, and alpha power. Note that there was a reduction of E/I in the right hemisphere according to Exponent and DFA-based E/I. The effects of iTBS on the contralateral hemisphere have never been mapped. The TMS studies performed on M1 contralateral to the site of stimulation have provided conflicting results (V. Di Lazzaro et al. 2008; 2011). Alpha power was also higher in the right hemisphere in this respect.

## Discussion

This pilot study demonstrated for the first time that EEG-based E/I estimates can mirror excitability changes produced by iTBS on the motor cortex as evaluated by TMS. We also considered for the first time multiple EEG-based E/I estimates, demonstrating that the majority of them exhibit expected modulations due to excitability tuning via iTBS.

### Are changes in excitability due to iTBS linked to changes in E/I estimated with EEG?

Previous studies demonstrated that iTBS induces an excitability increase in the stimulated M1 and a less consistent excitability decrease in the contralateral M1 (V. Di Lazzaro et al. 2011; Huang et al. 2005; Suppa et al. 2016; 2008). Previous evidence also suggests that the increase in excitability depends on the modulation of the E/I, through mechanisms that are specific to this neuromodulation technique (V. Di Lazzaro et al. 2008; Cárdenas-Morales et al. 2010). In detail, iTBS increases the late I-waves recorded at the cervical spine, suggesting selective influence on cortical interneurons producing the later components of the corticospinal volleys (V. Di Lazzaro et al. 2010). Single pulse TMS of the motor cortex interacts with neurons spanning all cortical layers, recruiting both excitatory and inhibitory cortical circuits (Vincenzo Di Lazzaro and Ziemann 2013). As such, spTMS MEPs are to be considered a rough measure of cortical excitability and are unable to disentangle the effects of iTBS on specific intracortical circuits. Furthermore, while iTBS mechanisms of action and effects have been characterized for M1 (V. Di Lazzaro et al. 2010; Suppa et al. 2008; V. Di Lazzaro et al. 2008), it is unclear whether they also apply to other cerebral cortices, where the cytoarchitecture is different. Therefore, it is not surprising that a TMS-EEG study on the effects of dorso-lateral prefrontal cortex iTBS did not appreciate remarkable effects on cortical excitability as the increase of global mean field power, but only of E/I (Desforges et al. 2022).

These are essentially the reasons why the relationship between excitability and E/I was tested for the left M1, where iTBS was applied.

Among all measures tested here, only the Exc_ind showed a correlation with changes in excitability probed by TMS (Figure 2). Meisel and collaborators developed this approach in 2015 as ongoing fluctuations of gamma phase synchrony as a proxy for intrinsic excitability (Meisel et al. 2015). In their studies, the global phase synchronization correlated with the amplitude of the cortical response following cortical stimulation, and with the presumed changes of excitability linked to changes in antiseizure medications (Ricci et al. 2021) and awake time (Meisel et al. 2015). Our pilot study extends previous findings in two relevant ways. Firstly, we demonstrate that this index is sensitive to enduring changes of excitability linked to neuromodulation and scales well with traditional excitability assessment via non-invasive TMS and recording of peripheral MEPs. Secondly, we provide insight into the spatial resolution of this measure. As the Exc_ind requires considering the phase synchrony across signals originating from spatially extended regions, it has an inherent limitation in terms of spatial resolution. In other words, it cannot be computed sensor-wise, or voxel-wise, or vertex-wise. Previously, the phase synchrony has therefore been mostly treated as a global property, reflecting global excitability (Meisel et al. 2016; 2015; Manzouri et al. 2021). We mapped excitability changes due to auditory stimulation by applying the Exc_ind on spatially resolved time-series extracted from cortical reconstruction of 275 sensors MEG data and considering, in the same way as in this study, the Desikan-Killiani atlas (Schuler et al. 2023). The present work suggests that the Exc_ind has at least Desikan-Killiany spatial resolution, but further studies are needed to determine if a higher resolution could be achieved. We relied on the Desikan-Killiany atlas for all measures as EEG recordings were performed with 32 electrodes only.

Nevertheless, the other measures of E/I are based on analysis of single time-series and can ensure higher spatial resolution, which is extremely important to achieve a non-invasive whole brain mapping of cortical excitability, especially in clinical contexts.

It remains to be unraveled whether the lack of significant relationship between changes in cortical excitability and E/I for other E/I estimates could be due to the lower sensitivity of these measures, the small sample size, or simply because other measures selectively assess particular aspects of E/I, which weight less in the global determination of M1 excitability.

### Do E/I estimates exhibit an expected modulation based on the known effects of iTBS?

Because of the known effects of iTBS (see above), it was reasonable to expect that EEG-derived E/I estimates would exhibit increases in excitability especially in motor regions of the left hemisphere. All E/I estimates but DFA showed the expected pattern (Figure 3). Differences across E/I estimates may be due to several reasons, intrinsic variability of the E/I estimate and sensitivity to different aspects of E/I.

The study of the exponent of the power spectrum as E/I estimate was motivated by the need to map E/I non-invasively and with high spatio-temporal resolution (Gao, Peterson, and Voytek 2017; Wilson, Da Silva Castanheira, and Baillet 2022). The validation of this index is based on the definition of E/I as the property determining the ongoing fluctuations of local field potentials. While some specificity for cortical layers has been demonstrated (Gao, Peterson, and Voytek 2017), when applied to electromagnetic signals recorded non-invasively with EEG or magnetoencephalography, this index likely reflects E/I across all cortical layers and how it influences the fluctuations of post-synaptic excitatory and inhibitory potentials. In our study, the exponent-based E/I showed an expected increase around the left M1 and an expected decrease in the contralateral hemisphere. Nevertheless, no significant correlation with excitability as measured by MEPs was found.

The entropy and the DFA-based E/I estimations are measures based on the temporal regularity of the signal. Their validation as measures of E/I has been largely performed in relationship to changes in consciousness and effects of medications (Sarasso et al. 2015; Wei et al. 2013; Liang et al. 2015).

Therefore, entropy and DFA-based E/I should be considered as broad and rather unspecific measures of E/I, and the mechanisms sustaining their link to excitability should be better ascertained. Here we demonstrated that sample entropy exhibited an expected pattern, whereas DFA-based E/I did not.

Alpha activity has been related to excitability patterns for a long time (Pellegrino et al. 2021). Here a role in conditioning the excitability is attributed to oscillations in specific frequency bands. This is therefore at the opposite of the exponent of the power spectrum, where E/I is inferred based on the spectral content of ‘background’ activity, after the removal of oscillations (Donoghue et al. 2020; Romei et al. 2008). Higher alpha power in M1 at the time of stimulation corresponds to lower excitability, as probed by online TMS (Sauseng et al. 2009). Here we show that alpha does indeed capture expected E/I changes induced by iTBS (Figure 3). Alpha oscillations are generated by an interplay between cortex and thalamus (Lopes da Silva 2013), and their specificity for E/I components is difficult to ascertain. Interestingly, three of five measures (Exc_ind, sample entropy and alpha power) showed a larger E/I increase in the regions contralateral to the iTBS target, than in the left M1. This is in line with recent TMS-fMRI findings (Tik et al. 2023) and might reflect some compensatory mechanisms.

Overall, E/I estimates exhibited an expected modulation based on the known effects of iTBS, but also showed differences between each other, underlining that they could capture complementary aspects of the E/I.

## Limitations

The main limitation of this study is the small sample size, which could be not sensitive enough to detect E/I modulations. In our cohort all participants showed an iTBS-induced increase in excitability, suggesting that at times MEP amplitude can be sensitive enough to appreciate variations up to single subject level. Conversely, there are no previous studies guiding on the sensitivity of E/I estimates.

Another limitation is that we considered the effects of iTBS only. Other neuromodulatory protocols, such as inhibitory and excitatory repetitive TMS, paired associative stimulation, transcranial direct current stimulation (tDCS), do induce changes in brain excitability through the modulation of other intracortical circuits. In other words, our experimental design may not generalize to all excitability changes and different E/I estimates could be sensitive to modulation of different intracortical circuits. The third limitation of the study is the low number of EEG channels. Nevertheless, in this pilot study we mostly focused on activity reconstructed in the left primary motor cortex, corresponding to the precentral gyrus of the Desikan-Killiani atlas. Here we have sufficient coverage and sensitivity and confidence in the accuracy of the reconstruction.

The fourth limitation is that not all EEG/MEG potential E/I estimates have been considered in this study (Pellegrino et al. 2017; Tecchio et al. 2018).

## Conclusions

EEG-based E/I estimates depict some degree of expected modulation linked to iTBS stimulation. Remarkable differences were found across E/I estimates, possibly due to variable sensitivity to different aspects of E/I balance and activity of intracortical circuits.

Only the so-called Exc_ind, based on the spatial phase synchrony of the high-gamma band, was a reliable proxy of cortical excitability modulation linked to iTBS. Overall, the estimation of cortical excitability from EEG-derived E/I index appears feasible. Further studies are needed for further characterization of the relationship between excitability and E/I balance and to translate the full potential of E/I estimates as a tool to map cortical excitability.

## Acknowledgments

GP is funded by a starting grant from the Department of Clinical Neurological Sciences, Schulich School of Medicine and Dentistry, Western University, London, Ontario.

